# Human Bone Marrow Adipose Tissue is a Hematopoietic Niche for Leptin-Driven Monopoiesis

**DOI:** 10.1101/2023.08.29.555167

**Authors:** Zinger Yang Loureiro, Amruta Samant, Anand Desai, Tiffany DeSouza, Haley Cirka, Mai Ceesay, David Kostyra, Shannon Joyce, Lyne Khair, Javier Solivan-Rivera, Rachel Ziegler, Nathalia Ketelut Carneiro, Linus T Tsai, Michael Brehm, Louis M Messina, Katherine A Fitzgerald, Evan D Rosen, Silvia Corvera, Tammy T Nguyen

## Abstract

During aging, adipose tissue within the bone marrow expands while the trabecular red marrow contracts. The impact of these changes on blood cell formation remains unclear. To address this question, we performed single-cell and single-nuclei transcriptomic analysis on adipose-rich yellow bone marrow (BMY) and adipose-poor trabecular red marrow (BMR) from human subjects undergoing lower limb amputations. Surprisingly, we discovered two distinct hematopoietic niches, in which BMY contains a higher number of monocytes and progenitor cells expressing genes associated with inflammation. To further investigate these niches, we developed an in-vitro organoid system that maintains features of the human bone marrow. We find cells from BMY are distinct in their expression of the leptin receptor, and respond to leptin stimulation with enhanced proliferation, leading to increased monocyte production. These findings suggest that the age-associated expansion of bone marrow adipose tissue drives a pro-inflammatory state by stimulating monocyte production from a spatially distinct, leptin-responsive hematopoietic stem/progenitor cell population.

**Significance:** This study reveals that adipose tissue within the human bone marrow is a niche for hematopoietic stem and progenitor cells that can give rise to pro-inflammatory monocytes through leptin signaling. Expansion of bone marrow adipose tissue with age and stress may thus underlie inflammageing.

## Introduction

Ageing and chronic metabolic stress are accompanied by a persistent low-grade inflammatory state known as inflammageing (1–3). Ageing and metabolic stress are also accompanied by the progressive expansion of the adipocyte-rich yellow bone marrow (BMY) (4–10). The yellow bone marrow first appears in human distal long bones by the age of seven and gradually expands, confining the trabecular red marrow (BMR) to the more proximal bone epiphyses (11, 12). The implications of these changes on hematopoiesis remain poorly understood largely due to lack of experimental platforms that model the human bone marrow. In addition, the study of adipose tissue is hampered by the fragility of adipocytes, which are large, lipid laden terminally differentiated cells.

Single-nuclei transcriptomics overcomes the limitations of single-cell transcriptomic platforms to analyze the cells comprising adipose tissues. Here, we leveraged single-nuclei transcriptomics to define cell types comprising the BMY and BMR of human adult long bones. To obtain insights on the functional roles of cells from BMY and BMR, we developed a 3D culture system (13–15) to generate organoids from the BMY and BMR of amputated long bones of adult human subjects. Bulk and single-cell RNA-seq (scRNA-seq) analyses of cultured BMY and BMR organoids closely matched the composition of their respective niches, as defined by single-nucleus RNA-seq (snRNA-seq). This validation underscores the capability of this platform to model human bone marrow, enabling experimental manipulation and functional investigation.

Transcriptomic analysis of BMY organoids confirm the finding that progenitor cells in this niche are enriched in genes associated with inflammation and monocyte development compared to cells derived from BMR. The increase in inflammatory gene profiles in cells expanded from BMY correlate with elevated leptin receptor (LEPR) expression, particularly in hematopoietic stem/progenitor cells (HSPCs) and monocytes. Functionally, addition of exogenous leptin to HSPCs derived from BMY enhances monocyte proliferation, resulting in monopoiesis and increased monocyte inflammatory cytokine secretion. LEPR^+^ progenitor cells also exhibit increased expression of colony-stimulating factor 1 (CSF1), known to stimulate monocyte differentiation and proliferation. These findings identify human bone marrow adipose tissue as a specialized niche enriched with multipotent LEPR^+^ mesenchymal stem/progenitor cell (MSPC) and HSPCs that support monopoiesis through leptin signaling. These findings suggest a mechanism linking expansion of bone marrow adipose tissue to pro-inflammatory hematopoiesis during ageing.

## Results

### Cellular composition of bone marrow depots assessed by single-nuclei RNASeq

We first sought to explore the distinguishing features of adipose tissue located within the diaphysis of the adult long bone (BMY) compared to subcutaneous adipose tissue (SAT), and to the adipose-poor tissue within the epiphysis (BMR) (Figure 1A and B). Adipose tissue harvested from SAT and BMY is predominantly occupied by unilocular adipocytes, a defining feature of white adipose tissue, whereas BMR has few adipocytes. Small, dense nucleated cells are observed in BMR, and between adipocytes in BMY, but not in SAT (Figure 1C). To study the cell composition of these three depots, we profiled 4,104 nuclei from SAT, 2,033 nuclei from BMY, and 3,136 nuclei from BMR tissue using snRNA-seq (Figure 1D, Supplemental Figure 1A). Using the Louvain clustering algorithm and Uniform Manifold Approximation and Projection (UMAP) plot, we identified 17 unique clusters among the three tissue depots that could be annotated based on established immune cell and subcutaneous adipose progenitor reference databases (Figure 1E, Supplemental Figure 1, B-C). Mature and progenitor cell types were identified from all three tissue depots at varying abundance (Figure 1F, Supplement Figure 1D). Compared to BMY, SAT had a high proportion of adipose mesenchymal progenitor cells (SAT progenitors), and higher abundance of smooth muscle cells, pericytes, and immune cells (Figure 1F). Interestingly, BMY had a higher proportion of mature adipocytes compared to SAT, mostly due to lower proportion of endothelial cells and SAT progenitors (Figure 1F). BMY had a greater number of erythroid progenitor and plasma cells while BMR had a greater number of osteoprogenitors and endothelial cells (Figure 1F). Bone marrow progenitors were negligible in SAT, but present in BMY at a greater percentage than BMR (Figure 1G, Supplemental Figure 1E). These results reveal that BMY is an abundant niche for specific subtypes of bone marrow progenitor cells. Indeed, further analysis indicate that bone marrow progenitor cells from BMY and BMR tissues differ substantially, as evidenced by their clustering into two separate groups in the UMAP projection (Figure 1H). These results suggest that BMY and BMR represent distinct hematopoietic niches. To further examine the immune cell type differences between these two bone marrow tissue depots, the immune cell cluster from Figure 1E was subclustered to identify specific immune cell types (Figure 1I). Interestingly, these results reveal that BMY is an abundant niche for monocytes whereas BMR is an abundant niche for T and NK cells (Figure 1J).

**Figure 1.**
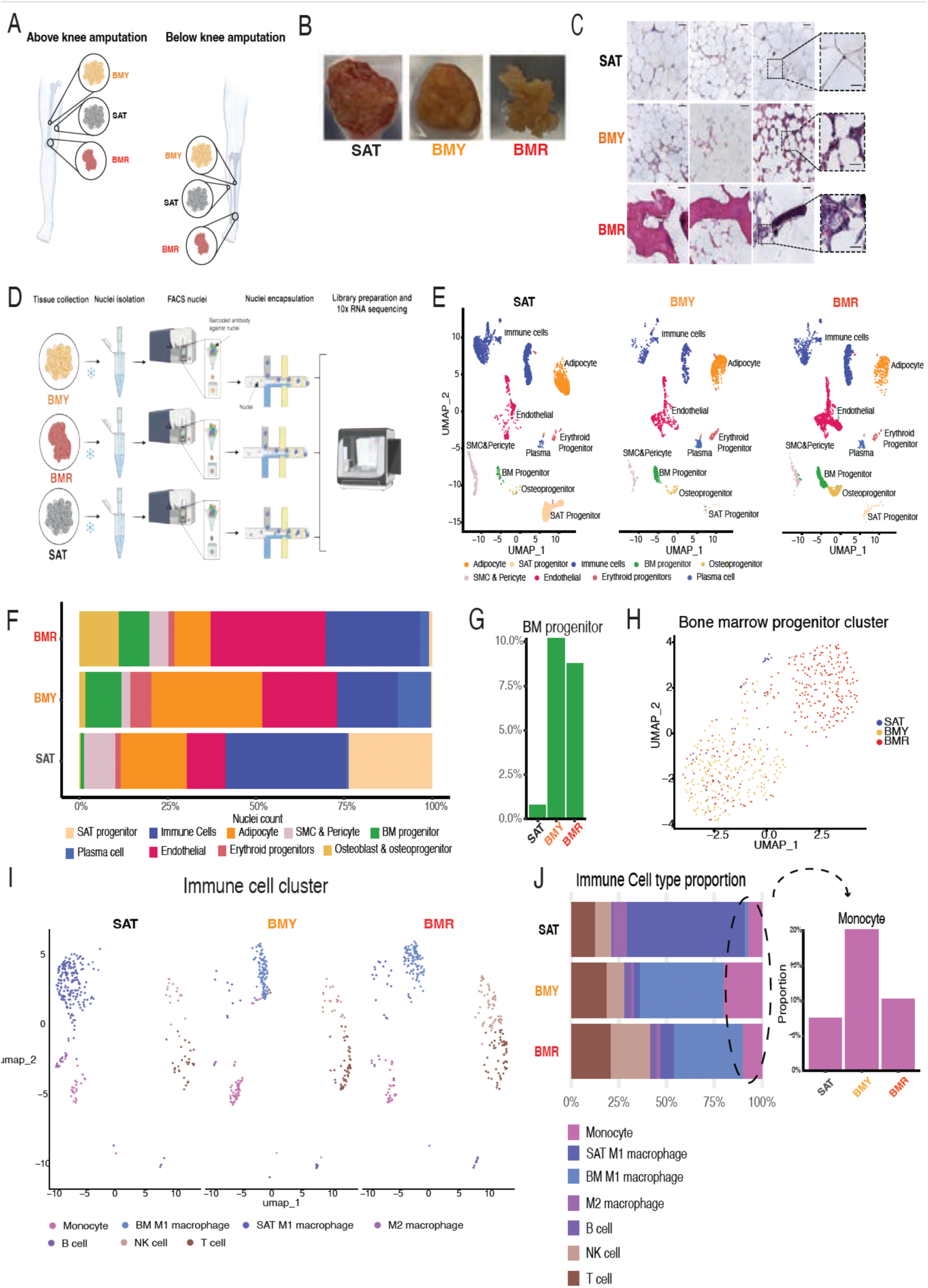
Features of adult human bone marrow. (A) Schematic of lower extremity bone with anatomic location of SAT, BMY, and BMR during above knee or below knee amputation. (B) Representative gross images of tissue depot. (C) Representative hematoxylin and eosin image of SAT, BMY, and BMR tissue harvested from three donors. Scale bar 200 μm. (D) Schematic of whole tissue nuclei extraction and isolation for snRNA Seq. (E) Dimensional reduction plot of snRNA Seq data using UMAP of SAT, BMY, and BMR tissue depots with cell clusters annotated to immune cell and adipocyte progenitor reference map (n = 1 human donor). (F) Composition of the percentage of each cell type nuclei over total nuclei for each tissue depot. (G) Bar graph of bone marrow (BM) progenitor percentage over total nuclei for each tissue depot. (H) UMAP visualization of bone marrow progenitor cells derived from SAT, BMY, and BMR tissues based on snRNA Seq, (n = 1 human donor). (I) UMAP visualization of immunce cell subclsuter derived from SAT, BMY, and BMR. (J) Bar graph of immune cell type percentage over 282 nuclei for each tissue depot, (n = 1 human donor).

### Human bone marrow organoids maintain cellular features of original tissue

To explore the functional roles of cells in each marrow compartment, we adopted a 3D culture system to develop tissue organoids (14). Small fragments of SAT, BMY, and BMR tissue were embedded in Matrigel and cultured in the presence of angiogenic growth factors (13, 14). Cells sprouted from all SAT, BMY, and BMR organoids collected from seven different donors (Figure 2A, Supplemental Figure 2A). After 14 days in culture, cells were recovered using dispase and collagenase Type I. A significantly higher number of cells were recovered from SAT organoids in comparison to those derived from BMY and BMR, possibly reflecting the higher proportion of mesenchymal progenitor cells in that depot (Figure 2B).

**Figure 2.**
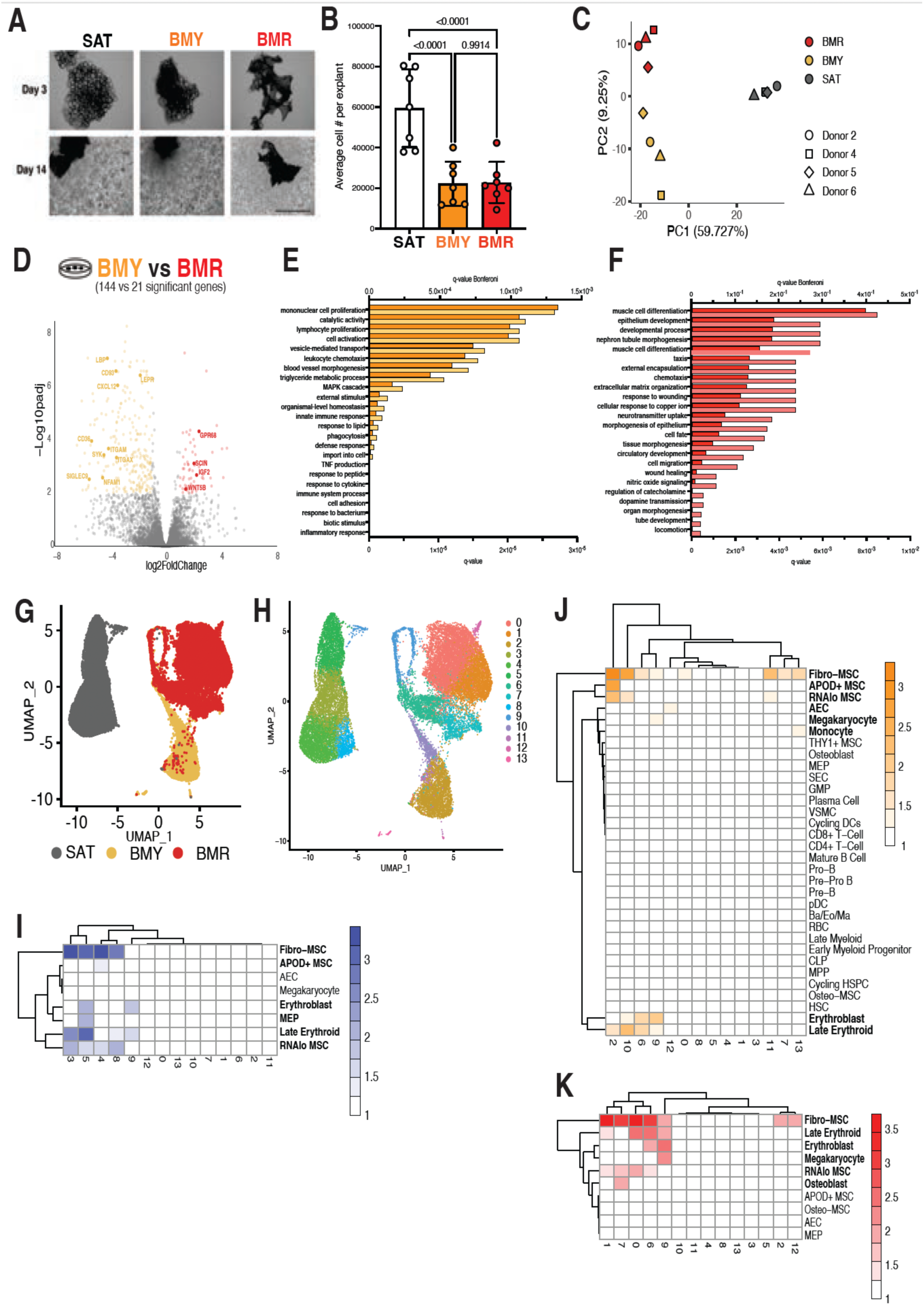
Human bone marrow adipose tissue progenitor cells are enriched for immune cell and inflammation pathways. (A) Representative images of SAT, BMY, and BMR tissue explant in 3D organoid culture system after 3 days and 14 days in culture under pro-angiogenic proliferation conditions. Scale bar 100 μm. (B) Average number of *de novo* cell expansion per 1 mm^3^ tissue explant organoid culture per tissue depot (n = 7 donors, bars = mean, error bars = SD, exact p-value indicated, p-value by two-way ANOVA). (C) Principal component analysis (PCA) was performed on differentially expressed gene between SAT, BMY and BMR (n = 4 human donors). (D) Volcano plot of differentially expressed genes between BMY in comparison to BMR. (E) Pathway enrichment analysis of differentially expressed genes in organoid cells derived from BMY in comparison to BMR cultured in mesenchymal stimulating media, q-Bonferroni value cut off < 0.05 and fold change > 4. (F) Pathway enrichment analysis of differentially expressed genes in organoid cells from BMR in comparison to BMY, q-Bonferroni value cut off < 0.05 and fold change > 4. (G) UMAP of SAT, BMY, and BMR organoid progenitors by tissue depot from scRNA Seq, (n = 1 human donor). (H) UMAP of SAT, BMY, and BMR organoid progenitor clusters based on 2000 gene features from scRNA Seq, (n = 1 human donor). (I) Heatmap of the assignment score for each UMAP cluster (column) and cell type label (row) from SAT based on Bandyopadhyay *et al.* (J) Heatmap of the assignment score for each UMAP cluster (column) and cell type label (row) from BMY based on Bandyopadhyay *et al.* (K) Heatmap of the assignment score for each UMAP cluster (column) and cell type label (row) from BMR based on Bandyopadhyay *et al*.

To compare the in vitro generated organoids with their tissue of origin, we first conducted bulk-RNA seq on samples from four independent subjects. Despite the diversity of the patient cohort sample, varying in sex, age, race, and co-morbidities (Supplemental Figure 2B), principal component analysis (PCA) of the organoid transcriptomes revealed the greatest variance corresponded to their depot of origin, indicating the maintenance of tissue identity despite prolonged culture (Figure 2C). Cells derived from BMY were more spaced in PC2, suggesting higher sensitivity to donor-specific variables.

DESeq analysis comparing BMY with BMR organoids (Figure 2D) revealed an enrichment of genes involved in inflammatory cell response pathways in BMY, and enrichment of genes associated with tissue development in BMR (Figure 2, E and F). These features are concordant with findings by snRNA-seq which show different transcriptomic profiles of progenitor cells from the two niches, with higher abundance of monocytes in BMY (Figure 1, H and J). Furthermore, we compared bulk RNA-seq of BMY, BMR, SAT organoids with bulk RNA-seq of freshly excised tissue from two separate donors. We found similar functional enrichments between freshly excised tissue and organoid cultures, with enhanced immune and hematopoietic regulatory pathways in BMY and enhanced bone developmental pathways in BMR (Supplemental Figure 2, C-F) when compared to SAT. The concordance between single-nuclei transcriptomic profiles from BMY and BMR tissues, and bulk RNA-Seq data from their corresponding organoids, demonstrates the validity of this culture system as a translational model for studying the functional features of the human bone marrow.

### Features of progenitor cells from SAT, BMY, and BMR organoids

To further compare the features of cells from SAT, BMY, and BMR organoids, we performed single-cell RNA-seq (scRNA-seq). After filtering out cells with low total sequencing count, high mitochondrial gene expression, and poor nuclear cell lysis, 8,688 cells derived from SAT, 3,974 cells derived from BMY, and 12,307 cells derived from BMR remained and were used for subsequent data analysis (Supplementa1 Figure 2, G-H). As seen in bulk RNASeq data, clustering of single cells was defined by tissue type, further supporting the finding the identity of each bone marrow niche is maintained during 3D culture. Cells from SAT organoids were most distinct, while cells from BMY and BMR clustered closer to each other (Figure 2G).

When analyzing for 2000 gene features within this scRNA-seq dataset, we observed 14 unique cell clusters from SAT, BMY, and BMR organoids (Figure 2H). We mapped these clusters to an existing human bone marrow scRNA-seq data set (16). Interestingly, of the six MSPC subtypes defined by Bandyopadhyay et al., fibro-MSCs, which are a more primitive cell type, were more abundant in all three organoids compared to the more differentiated MSPC subtypes (adipo-MSCs and osteo-MSCs), suggesting that organoid cultures promote proliferation of more naive human MSPCs (Figure 2, I-K). Cell clusters within SAT (Figure 2I) are primarily MSC and erythroid origin, whereas BMY and BMR cell clusters express other progenitor cell types (Figure 2J-K). Strikingly, we found that BMY cluster 13 is enriched in monocyte gene expression, and is not present in BMR or SAT (Figure 2J). In contrast, BMR cluster 7 is enriched in osteoblast gene expression (Figure 2K). The high enrichment of monocytes observed in BMY organoids and tissue (Figure 2I, Figure 1J) may explain the enrichment of inflammatory immune pathway genes in this niche observed in bulk RNA-seq (Figure 2E). Additionally, pathway analysis of progenitor cells using snRNA-seq from tissue and scRNA-seq from organoids revealed enrichment of myeloid supportive genes in BMY from both sources (Supplemental Figure 2I). Overall, the data demonstrate that the organoids accurately reflect in vivo findings, and underscore BMY as a bone marrow depot enriched in myeloid and monocytes at various developmental stages.

### Expansion of HSPCs in human bone marrow adipose tissue

To further explore whether hematopoiesis occurs in BMY, we supplemented the organoid culture media with factors known to support hematopoietic stem cell expansion, including stem cell factor (SCF), interleukin-6 (IL-6), thyroperoxidase (TPO), Fms-like tyrosine kinase 3 ligand (Flt3), pyrimido-indole derivative (UM171 or UM729), and StemRegenin (SR-1) (17–19). The addition of this cocktail (HSC media) increased the appearance of small round cells in BMY and BMR organoids but not in SAT (Figure 3A). While the addition of HSC media slightly decreased total cell recovery (Figure 3B, Supplemental Figure 3, A and B), it enhanced expression of hematopoietic gene regulatory pathways in BMY and BMR (Figure 3, C and D).

**Figure 3.**
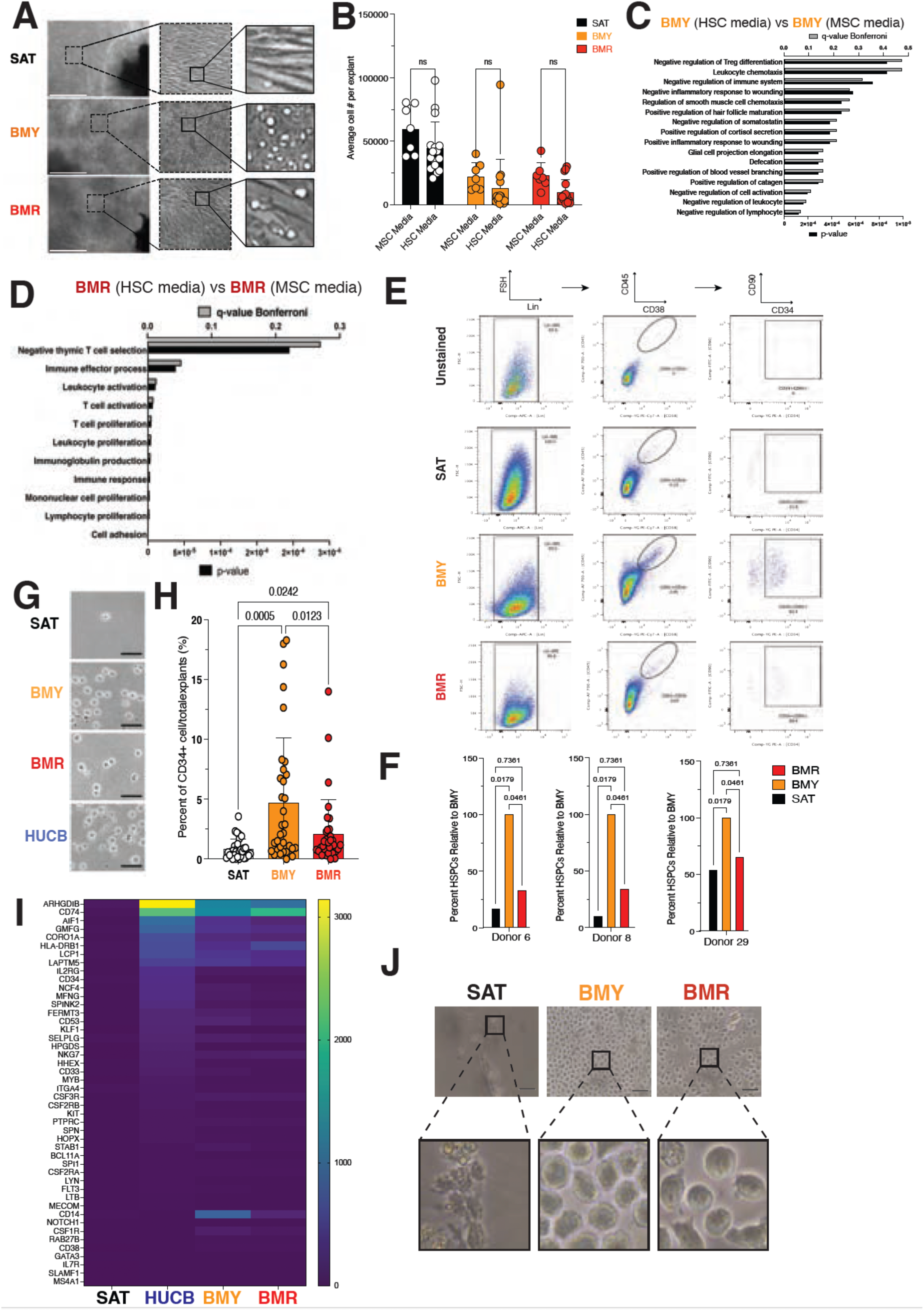
Human hematopoietic stem progenitor cell derived from bone marrow adipose tissue perivascular niche. (A) Representative images of SAT, BMY, and BMR tissue explants in 3D organoid culture system after 14 days in culture under hematopoietic proliferation and pro-angiogenic conditions. Scare bar 1000 μm. (B) Number of cells per 1 mm^3^ tissue explant after 14 days in mesenchymal (MSC) or hematopoietic (HSC) proliferative media conditions (n = 7 donors and n = 17 donors respectively, bars = mean, error bars = SD, as indicated by two-way ANOVA). (C) Pathway enrichment analysis of differentially expressed genes in BMY cultured in HSC expansion media in comparison to BMY cultured in MSC media, q-Bonferroni value cut off <0.05 and fold change >4. (D) Pathway enrichment analysis of differentially expressed genes in BMR cultured in HSC expansion media in comparison to BMR cultured in MSC media, q-Bonferroni value cut off <0.05 and fold change >4. (E) Representative flow cytometry analysis of organoid cells from SAT, BMY, and BMR explants. HSPC population box represents cells with canonical HSPC cell surface markers, Lin^-^ CD45^+^ CD38^-^ CD34^+^ CD90^+^. (F) Quantification of cell population positive for canonical HSPC cell surface markers, Lin^-^ CD45^+^ CD38^-^ (n = 3 donors per tissue depot, bars = mean, error bars = SD, exact p-value indicated by two-way ANOVA). (G) Representative images of CD34^+^ cells magnet bead purified from SAT, BMY, and BMR organoids and HUCB after dispase, collagenase, and trypsin digestion from Matrigel. Scale bar = 100 μm. (H) Quantification of the percentage of CD34^+^ cells purified from the total number of digested cells derived from SAT, BMY, and BMR 3D organoid culture (n = 35 donors per tissue depot, bars = mean, error bars = SD, exact p-value indicated by one-way ANOVA). (I) Bulk RNA-seq of marker hematopoietic genes expressed in CD34^+^ cells enriched from HUCB, SAT, BMY, and BMR 3D organoid culture (n = 1 SAT donor, 1 HUCB donor, 3 BMY donors, 3 BMR donors). (J) Representative 40x magnification images of CD34^+^ cells from SAT and BMY organoids cultured at day 18. Scale bar = 50um.

To determine whether cells expanding in SAT, BMY and BMR organoids in response to HSC media correspond to HSPCs, we probed for cell surface markers Lin^-^, CD45^low^, CD38^-^, CD34^+^, CD90^+^ (Figure 3, E and F, Supplemental Figure 3C). Cells with HSPC cell surface markers were detected in BMY and BMR, but not in SAT organoids. We observed a relative higher percentage of HSPCs (Lin^-^, CD45^low^, CD38^-^, CD34^+^, CD90^+^) in BMY when compared to SAT and BMR from three independent donors (Figure 3F). To test whether cells with HSPC cell surface markers have hematopoietic potential, we further isolated CD34^+^ cells using an enrichment bead purification method. CD34^+^ cells recovered from BMY and BMR were similar in size and shape to those derived from human umbilical cord blood (HUCB) and virtually no cells were recovered from SAT (Figure 3G). There was a trend for an increased number of CD34^+^ cells expanded from more proximal bone marrow compared to distal bone marrow for both BMY and BMR (Supplemental Figure 3, D and E). Interestingly, the mean number of CD34^+^ cells derived from BMY was greater to that from BMR organoids (Figure 3H). Since CD34^+^ cells may represent a mixed population of HSPCs and non-HSPCs, we probed for HSPC specific genes and found several to be expressed in CD34^+^ cells derived from BMY, BMR, and HUCB, but not from SAT (Figure 3I). We also observed increased CD14 expression, canonical gene marker for monocytes, from CD34^+^ cells derived from BMY organoids, corroborating with previous snRNA-seq from tissue and scRNA-seq from organoid data supporting monocyte enrichment in BMY (Figure 3I). Moreover, the few CD34^+^ cells derived from SAT organoids did not expand when cultured in HSPC expansion media, while those from BMY and BMR did (Figure 3J). These results indicate that HSPC can be expanded from BMY and BMR organoids.

To further characterize the HSPC subpopulations, we probed cells from organoids for common myeloid progenitor (CMP) cells (Lin^-^, CD34^+^, CD45RA^-^, CD38^+^), and common lymphoid progenitor (CLP) cells (Lin^-^, CD34^+^, CD45RA^+^, CD38^+^), using HUCB as a positive control. A negligible number of CMP and CLP cells expanded from SAT organoids, while HUCB was enriched with both (Figure 4, A and B). BMY organoids had more CMP cells compared to BMR (Figure 4, A and B, Supplemental Figure 4A).

**Figure 4.**
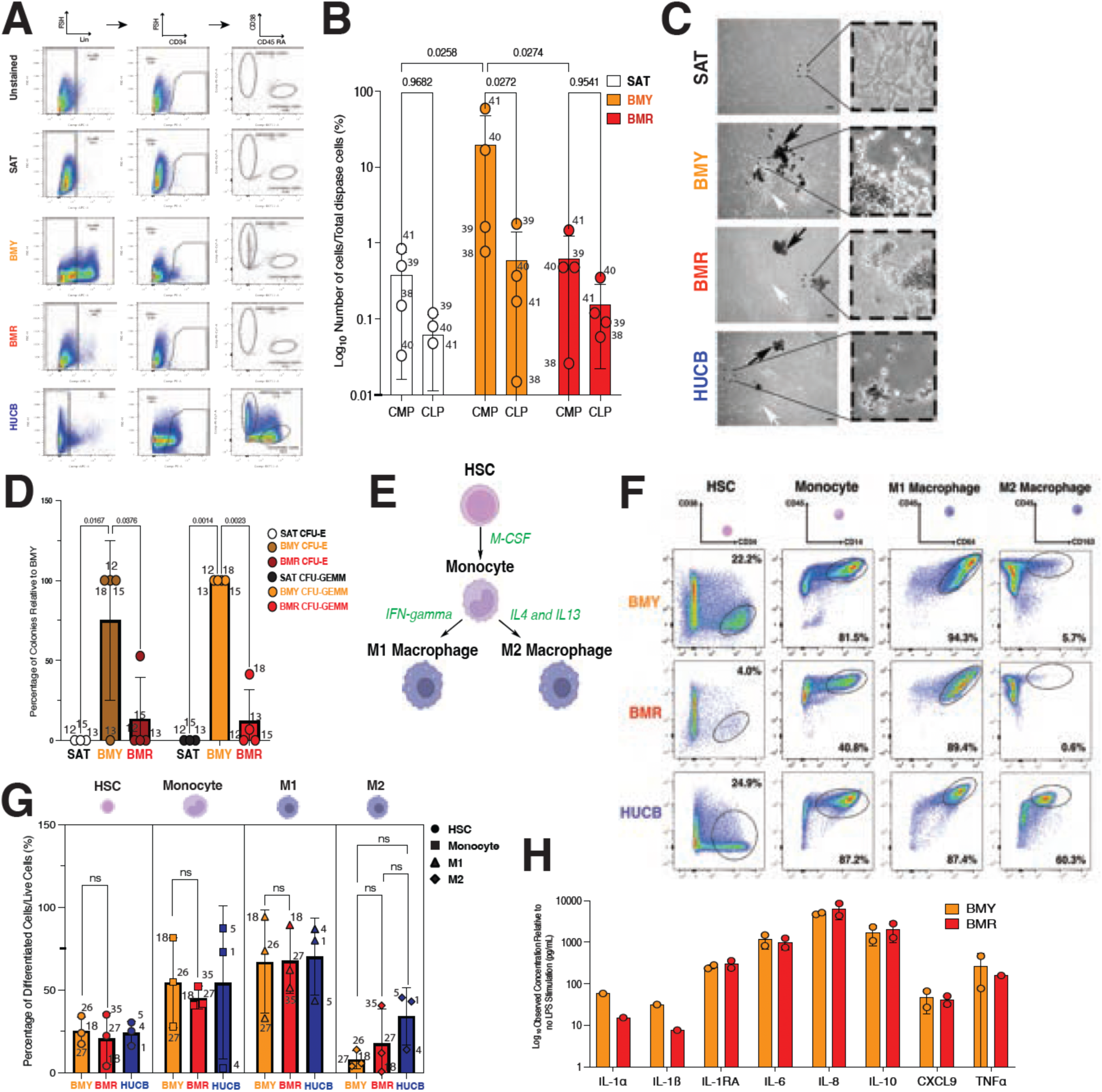
Bone marrow adipose tissue support myeloid progenitor cells. (A) Representative flow cytometry of common myeloid progenitor (CMP, Lin^-^, CD34^+^CD38^+^CD45RA^-^) and common lymphoid progenitor (CLP, Lin^-^, CD34^+^CD38^+^CD45RA^+^) cell surface markers expressed from SAT, BMY, and BMR organoids, and HUCB cultured in Matrigel for 14 days. (B) Quantification of cell population positive for CMP and CLP cell surface markers expressed from SAT, BMY, and BMR organoids (n = 4 donors per tissue depot, bars = mean, error bars = SD, exact p-value indicated by two-way ANOVA, donor sample indicated). (C) Representative images of colony forming units derived from CD34^+^ cells purified from SAT, BMY, and BMR tissue explants and HUCB. Black arrows indicate erythroid colonies. White arrows indicate myeloid colonies. Scale bar = 10 μm. (D) Quantification of colony forming units per 1000 seeded CD34^+^ cells. Indicated bars represent erythroid colonies (CFU-E) and granulocyte, erythrocyte, monocyte, macrophage (CFU-GEMM) colonies (n=4 donors, error bars = SD, exact p-value indication by two-way ANOVA, donor sample indicated). (E) Schematic for CD34^+^ differentiation into monocytes, M1 pro-inflammatory macrophages, and M2 anti-inflammatory macrophages. (F) Representative flow cytometry of HSPC (Lin^-^CD34^+^CD38^-^), monocyte (CD14^+^CD45^+^), M1 pro-inflammatory macrophages (CD63^+^CD45^+^), and M2 anti-inflammatory macrophages (CD163^+^CD45^+^ or CD206^+^CD45^+^) cell surface markers differentiated from BMY and BMR organoids and HUCB cultured in Matrigel for 14 days. (G) Quantification of cell population positive for indicated cell type based on cell surface markers expressed from BMY and BMR organoids and HUCB (n = 4 donors per tissue depot, bars = mean, error bars = SD, exact p-value indicated by two-way ANOVA, donor sample indicated, HUCB sample 1, 4, and 5). (H) Quantification of cytokine secretion after LPS stimulation from monocytes derived from BMY and BMR organoids, n = 2 donors per tissue depot (donor 36 and 40), bars = mean, error bars = SD.

In a colony forming unit assay, CD34^+^ cells from BMY, BMR and HUCB formed both erythroid, granulocyte, monocyte, and megakaryocyte colonies, whereas CD34^+^ cells from SAT formed neither (Figure 4C, Supplemental Figure 4, B and C). Moreover, BMY yielded a higher number of erythroid and granulocyte, monocyte, and megakaryocyte colonies per number of CD34^+^ cells compared to BMR (Figure 4D). These data demonstrate that functional HSPCs capable of myeloid differentiation proliferate in BMY and BMR organoids. The presence of an increased CMP cell population and more efficient myeloid cell differentiation observed in BMY relative to BMR reflects an enrichment for myeloid lineage differentiation within bone marrow adipose tissue.

We then explored whether CD34^+^ cells derived from BMY and BMR can differentiate into monocytes and macrophages. Monocyte and macrophage differentiation was observed in cells from both bone marrow organoid types and was comparable to that seen from HUCB (Figure 4, E-G, Supplemental Figure 4D). To probe for functional features of monocytes derived from either BMY or BMR we measured cytokine production after LPS stimulation (Figure 4H). We find similar, strong stimulation of cytokine production by monocytes differentiated from either BMY or BMR.

### Leptin stimulates monopoiesis in bone marrow adipose tissue

To explore the mechanisms underlying the increased CMP and monocyte production seen from BMY organoids, we evaluated for differential gene expression between BMY and BMR analyzed by scRNA-seq, and find the leptin receptor (*LEPR*) is the most differentially expressed gene from BMY compared to BMR (Supplemental Figure 5A). We then performed pseudobulk analyses of the transcriptomes of CD34^-^ and CD34^+^ cells to evaluate for LEPR expression. We find that LEPR expression is enriched in BMY among both CD34^+^ and CD34^-^ cell populations (Figure 5A). CD34^+^ cells represent 1.7% of the SAT, 0.94% of the BMY, and 0.48% of the BMR cell population (Supplemental Figure 5B). To test whether LEPR protein is expressed in CD34^+^ HSPCs, we performed FACS analysis probing for lineage negative cells expressing both LEPR and CD34^+^. These cells were more abundant among Lin^−^ cells originating from BMY (Figure 5, B and C).

**Figure 5.**
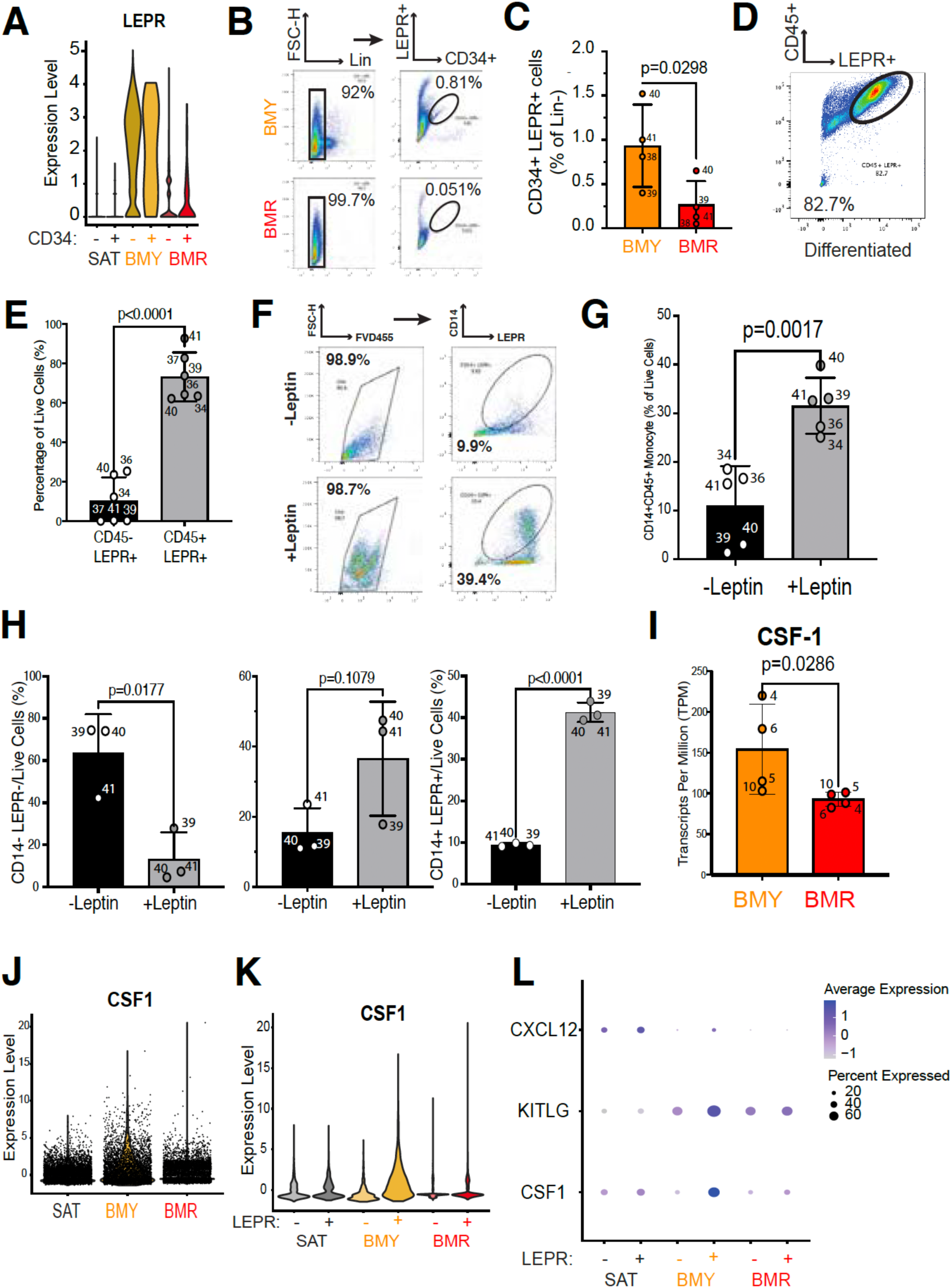
Leptin signaling enhances monocyte production. (A) Violin plot of *LEPR* expression variance in BMR, BMY, and SAT (n=1 donor). (B) Representative flow cytometry analysis of LEPR^+^ CD34^+^ cell population from total lineage negative cells cultured from 3D organoid culture. LEPR^+^CD34^+^ cell population percentage represents cells in oval. (C) Quantification of the percentage of LEPR^+^ CD34^+^ cells from total lineage negative cells from 3D organoid culture (n = 4 donors, error bars = SD, exact p-value indicated by Student’s t-test, donor sample indicated). (D) Representative flow cytometry analysis of. LEPR^+^CD45^+^ cell population cultured in M-CSF (differentiated) media. (E) Quantification of the percentage of live cultured cells expressing both LEPR^+^ CD45^-^ markers (undifferentiated) or LEPR^+^ CD45^+^ (differentiated) markers cultured in M-CSF media based on flow cytometry (n = 4 donors, error bars = SD, exact p-value indicated by Student’s t-test, donor sample indicated). (F) Representative flow cytometry analysis of CD14^+^ monocytes expressing LEPR^+^ with and without leptin stimulation. (G) Quantification of the percentage of CD14^+^ CD45^+^ monocytes with and without exogenous leptin based on flow cytometry (n = 5 donors, error bars = SD, exact p-value indicated based on Student’s t-test, donor sample indicated). (H) Quantification of the percentage of LEPR^-^CD14^-^, LEPR^+^CD14^-^, and LEPR^+^CD14^+^ cell population with and without exogenous leptin based on flow cytometry (n = 3 donors, error bars = SD, exact p-value is indicated based on Student’s t-test). (I) *CSF1* expression in organoid cells based on bulk RNA-seq. (n = 4 donors, error bars = SD, exact p-value indicated by Student’s t-test, donor sample indicated). (J) Violin plot of *CSF-1* expression variance in SAT, BMY, and BMR based on scRNA-seq (n = 1 donor). (K) Violin plot of *CSF1* expression in LEPR^-^ and LEPR^+^ cells purified from SAT, BMY, and BMR 3D organoid culture based on scRNA-seq. (n = 1 donor). (L) Dot plot of *CXCL12*, *KITLG* (*SCF*), and *CSF1* gene expression in LEPR^+^cells and LEPR^-^ cells derived from SAT, BMY, and BMR based on scRNA-seq (n = 1 donor).

To test whether LEPR expression is maintained during differentiation, we cultured CD34^+^ cells isolated from BMY organoids with or without M-CSF for 5 days to induce for monocyte differentiation. The majority of differentiated CD34^+^ cells displayed the mature immune cell surface marker CD45 and LEPR after induction with M-CSF (Figure 5D). We observed that 73% of live differentiated cells expressed both LEPR and CD45^+^, while LEPR expression was detected in only 10% of undifferentiated live CD45^-^ cells (Figure 5E). These data indicate that LEPR expression is maintained during monocyte differentiation of CD34^+^ cells from BMY.

To determine the functional role of the LEPR receptor, we cultured CD34^+^ cells derived from BMY organoids with M-CSF, either in the absence or presence of exogenous leptin, and measured the number of differentiated monocytes in the population using CD14. Leptin increased the number of CD14^+^ CD45^+^ monocytes and CD14^+^ cells expressing LEPR (Figure 5, F and G, Supplemental Figure 5, C and D). Moreover, the proportion of CD14^+^ LEPR^+^ cells in the culture increased from ∼10 to ∼40% of live cells while the proportion of CD14^-^ LEPR^-^ cells decreased in the presence of leptin, suggesting a positive myeloproliferative feedback loop (Figure 5H).

The data shown above indicates that monocyte production occurs in BMY and can be enhanced through direct leptin signaling. To examine whether LEPR^+^ non-hematopoietic progenitor cells may also play a role in supporting monocyte proliferation, we queried our data for key factors that support myelopoiesis. We found that cells from BMY organoids display higher expression of CSF1/M-CSF compared to BMR (Figure 5I), confirmed by pseudo-bulk analysis of scRNA-seq datasets (Figure 5J). Moreover, increased CSF1 expression was more pronounced in LEPR^+^ cells from BMY (Figure 5K). BMY LEPR^+^ cells also displayed enrichment of other ligands such as CXCL12 and c-kit ligand (KITLG or SCF) (Figure 5L). These findings suggest that BMY derived LEPR^+^ cells can sustain an HSPC population through SCF and CXCL12 secretion and monopoiesis through CSF1 signaling.

## Discussion

The inverse relationship between expansion of bone marrow adipose tissue and loss of trabecular red bone marrow during ageing and metabolic disease is much more pronounced in humans than mice (20). However, the physiological impact of this age-associated phenotype has been difficult to address given the paucity of methods to study the functional features of the human bone marrow adipose tissue. In this study, we developed methods to investigate the composition and functional properties of human bone marrow niches from human donors undergoing lower extremity amputations. The main finding in this study is that, compared to BMR, BMY is enriched for inflammatory immune cell pathways and monocytes suggesting that the BMY niche may be a preferential site for monopoiesis. Monocyte-enriched hematopoiesis is commonly observed during acute stress such as infection (21). Based on the characteristics of the study cohort, comprised of individuals undergoing surgery due to diverse and multiple conditions including infection, ischemia, and trauma, our findings may reflect the response of the human bone marrow to physiological stress.

The physiological role of the bone marrow adipose tissue has been difficult to elucidate, as the large volume and fragility of the abundant adipocytes within this niche preclude cell isolation in mice and human. Through snRNA-seq transcriptomic analysis, we found that BMY harbors an increased number of monocytes and progenitor cells expressing inflammatory gene signatures compared to BMR. To further explore the functional properties of the human BMY and BMR while preserving bone marrow tissue structure, we developed an in vitro organoid system that preserves key features of the human bone marrow. Bone marrow organoids scRNA-seq transcriptomics corroborated with several cell types and gene features identified in corresponding tissue transcriptomics, confirming the validity of this model to elucidate the physiological roles of these bone marrow depots. We further find that HSPCs are produced by both BMY and BMR organoids, despite vast differences among our donors’ demographic characteristics. HSPCs from both BMY and BMR give rise to common myeloid and lymphoid progenitors, erythroid and myeloid colony forming units, and differentiated monocyte and macrophage populations. Functionally, HSPCs derived from the BMY form erythroid and myeloid colonies more efficiently than those from BMR (Figure 4, C and D), and displayed a pronounced monopoiesis potential (Figures 1J, 2J, 3I).

The technology to expand progenitors that retain in vivo features of BMY allowed us to explore the cellular and molecular function of adipose tissue and human hematopoiesis. We found that the LEPR is differentially expressed in both HSPCs and MSPCs derived from BMY organoids (Figure 5A). LEPR^+^ cells have been identified in the perivascular niche of the bone marrow and found to be primarily expressed on murine bone marrow MSPCs for HSC maintenance through secretion of SCF/KITL and CXCL12 (22, 23). In addition to the expression of LEPR in MSPCs, CD34^+^ LEPR^+^ cells have recently been reported to represent a small subpopulation of long-term HSCs (LT-HSCs) with enhanced engraftment capacity in young mice. Interestingly, CD34^+^ LEPR^+^ increase in population size and lose engraftment efficiency with ageing (24). Our results indicate that CD34^+^ LEPR^+^ HSPCs are present in human and are more abundant in BMY (Figure 5, D and E). Furthermore, human differentiated CD34^+^ LEPR^+^ HSPCs were sensitive to leptin stimulation (Figure 5, F and G). Leptin also enhances the proliferation of LEPR^+^ monocytes derived from BMY organoids, indicating a positive myeloproliferative feedback loop initiated by leptin signaling (Figure 5H).

Leptin, a hormone secreted by adipocytes, may play a direct role in shaping hematopoietic activity within BMY. We found that human LEPR^+^ bone marrow progenitor cells derived from BMY have increased CSF1 expression (Figure 5, L and M) when compared to BMR. Two independent scRNA-seq studies in mice revealed increased CSF1 expression in LEPR^+^ cells relative to other progenitor cell types (25). In mice, conditional deletion of CSF1 leads to a reduction in the total number of monocytes in the animal (25). CSF1/M-CSF is continuously secreted under steady-state conditions and rises during acute stress and inflammation, triggering emergency myelopoiesis to supply the increased systemic demand for mature monocytes (21, 26–29). Our results indicate an increased expression of CSF1 in progenitor cells derived from BMY (Figure 5, I-L), which could contribute to the monocyte enrichment from this niche. It is possible that a conserved leptin signaling and CSF1/M-CSF expression may promote stress and/or age-related monopoiesis that is enhanced in human due to an increase in bone marrow adipose tissue abundance relative to mice. Further studies will be needded to reveal molecular relationships between leptin signaling in human bone marrow and downstream changes in hematopoiesis.

The differences in HSPC abundance between BMY and BMR organoids may reflect our culture conditions or the diverse clinical conditions of the study cohort, which includes individuals undergoing surgery for infection, ischemia, and trauma. To fully understand the molecular and functional distinctions between HSPCs in the BMY and BMR niches, larger cohort studies that are designed for donor demographic stratification will be necessary in the future. Despite the donor diversity within this study, our findings consistently show that the human BMY niche is enriched with monocytes and bone marrow progenitor cells that support monopoiesis, as demonstrated by both in vivo and in vitro organoid studies. BMY is unlikely the sole niche for human monopoiesis, however, the enrichment of monocytes in BMY tissue and organoids, along with the leptin-driven monocyte production, suggests that bone marrow adipose tissue plays an active role in shaping human hematopoiesis.

## Methods

### Sex as a Biological Variable

Male and female patients were included in this study; 65% of the cohort was male and 35% was female.

### Study Approval

Lower extremity subcutaneous adipose tissue, yellow bone marrow adipose tissue, and red marrow adipose tissue were collected from discarded tissue of patients undergoing lower extremity amputation, with approval from the University of Massachusetts Institutional Review Board (IRB H00020377).

### Data Availability

Sequencing results were submitted to GEO (accession number: GSE231377). Sequencing results and analysis scripts will be made publicly available upon publication of the manuscript.

### Statistical Analysis

Statistical analyses were performed using GraphPad Prism software version 10.0. For normal distributions, parametric Student’s *t* tests were used. In analyses where multiple groups were compared, a one-way or two-way parametric ANOVA was used with subsequent multiple comparison. Two-tailed statistical tests were always used. A *P* value <0.05 was considered significant.

### Code Availability

Only open-source software was used for this study, and no custom code packages were generated for data analysis. Please refer to the computing package citations.

### Histology

All samples were fixed in 4% paraformaldehyde at 4°C and thoroughly washed with PBS. Tissue sections (10 mm) were mounted on Superfrost Plus microscope slides (Fisher Scientific) and stained with hematoxylin and eosin. Tissue sections were imaged using ZEISS Axio Scan.Z1.

### Generation of 3D tissue explant co-culture system

Methods for the harvesting and the culture adipose tissue explants were previously published (15). Human tissues were donated from consented adult patients undergoing lower extremity amputation surgery at University of Massachusetts Medical Center and were subjected to harvesting within one to two hours following amputation. Explants of approximately 1cm^3^ in size were embedded in Matrigel Matrix (Cat# 356231, Corning) per 10 cm dish with EGM-2MV (Cat# CC-3156,CC-4147, Lonza) media supplementation. The excised tissue was embedded in Matrigel (180 explants/10 cm dish) or (30 explants/6-well dish) and cultured for 14-17 days as described before. Addition of 100ng/mL stem cell factor (SCF), 100ng/mL interleukin-6 (IL-6), 100ng/mL thyroperoxidase (TPO), 100ng/mL Fms-like tyrosine kinase 3 ligand (Flt3), 35mM pyrimido-indole derivative (UM171 or UM729), and 0.75 μM StemRegenin (SR-1) were added to culture to promote HSC expansion. After 12-17 days in Matrigel, the progenitors in explants were recovered using Dispase (Cat# 354235, Corning) for one hour followed by an additional 14-minute incubation of Trypsin-EDTA (Cat# 15400-054, Gibco) and Collagenase I (Cat# LS004197, Worthington).

### RNA extraction and quantitative PCR

Cells were placed in TRIzol (Invitrogen) and homogenized with Tissuelyser (Qiagen). Total RNA was reverse transcribed using the iScript™ cDNA Synthesis Kit (Cat# 1708891, Bio-Rad) per manufacturer’s protocol. Quantitative reverse-transcription PCRs were prepared with iQTM SYBR Green Supermix (Cat# 1708882, Bio-Rad) and were performed on a CFX Connect Real-Time PCR Detection System (Bio-Rad). Amplification of human-specific genes was accomplished using the following primers:

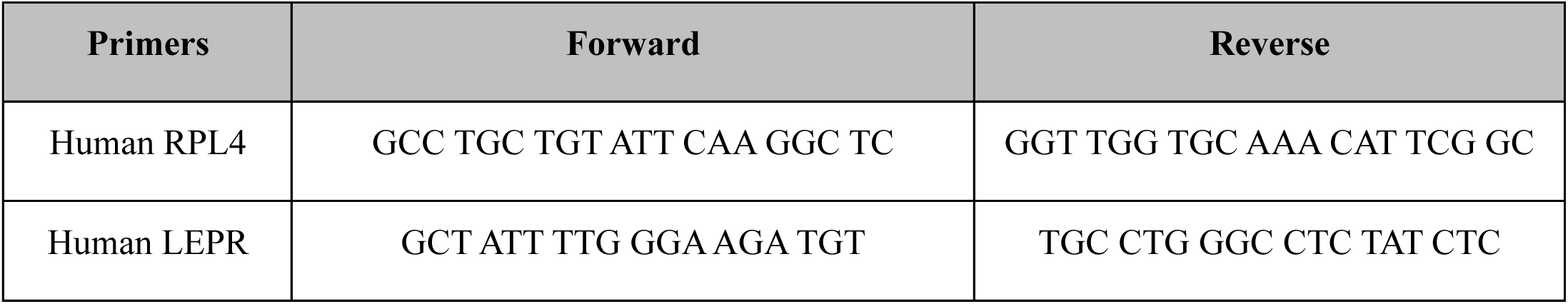

### Nuclei isolation and snRNA-Seq

Nuclei were prepared for snRNA-Seq using the Chromium™ Single Cell 3’ GEM Library & Gel Bead Kit v3.1 (Cat# 1000121, 10X Genomics) according to protocol described in Gulko, 2024 (30). Briefly, each flash-frozen adipose tissue sample was dissociated using a gentleMACS Dissociator (Cat # 1390-093-235, Miltenyi Biotec) and 2 mL TST buffer (0.03% Tween 20 (Cat # 1610781, Bio-Rad), 0.01% BSA (Cat # B8999S, New England Biolabs), 146 mM NaCl (Cat # AM9759, ThermoFisher), 1 mM CaCl_2_ (Cat # 97062-820, VWR), 21 mM MgCl_2_ (Cat # M1028, Sigma Aldrich), and 10 mM Tris-HCl pH 7.5 (Cat # 15567027, ThermoFisher) in Ultrapure water (Cat # M1028, ThermoFisher)) with 0.2 U µl^−1^ of Protector RNase Inhibitor (Cat # 3335402001, Sigma Aldrich), by running the program mr_adipose_01 3 times. Following incubation on ice for 10 min, lysate was passed through a 40 µm nylon filter, rinsed with 3 ml ST buffer (146 mM NaCl, 1 mM CaCl_2_, 21 mM MgCl_2_; 10 mM Tris-HCl pH 7.5), 0.2 U/µl RNase Inhibitor), and subsequently passed through a 20 µm filter to remove aggregates. Flow-through was centrifuged at 500*g* for 5 min at 4 °C with brake set to half-maximal. Following centrifugation, supernatant was removed, the nuclear pellets were washed with 5 ml Nuclei Resuspension Buffer (NRB) (PBS 7.4 (Cat # 10010023, ThermoFisher), 0.02% BSA, 0.2 U/µl RNase inhibitor) and centrifuged with identical settings before resuspension in 500 µl NRB. Nuclei for each sample were stained with one drop of NucBlue (Cat # R37605, ThermoFisher) and 2 ul of anti-nuclear hashtag antibody (Cat # 682205, 682207, 682209, BioLegend) and incubated for 45 minutes. Following incubation, the labeled nuclei were pelleted by centrifugation and resuspended in 500 uL NRB. The labeled nuclei from each tissue were then sorted (Beckman Coulter MoFlo Astrios EQ, 70 mm nozzle) into the same well and immediately loaded on the 10x Chromium controller (Cat # PN120270, 10X Genomics) according to the manufacturer’s protocol. 17334 nuclei from each sample (52002 nuclei total) were loaded in one channel of a Chromium Chip, and cDNA and gene expression libraries were generated according to the manufacturer’s instructions. cDNA and gene expression library fragment sizes were assessed with a DNA High Sensitivity Bioanalyzer Chip (Cat # 5067-4626, Agilent). cDNA and gene expression libraries were quantified using the Qubit dsDNA High Sensitivity assay kit (Cat # Q33231, ThermoFisher). The libraries were sequenced on the NovaSeq system (Illumina) by Azenta Life Science. The sequencing outputs were processed using the CellRanger software v3.1.0. Reads were mapped to human reference genome hg38. Data analysis was performed using Seurat v5.0.1 within R version 4.3.1 environment(31). Sequencing results will be made publicly available upon publication of the manuscript.

### Bulk RNA-seq

RNA was extracted from dispased cells using the TRIzol method. The cell-TRIzol mixtures were transferred to collection tubes and homogenized with Tissuelyser II (Qiagen). Chloroform was added in a 1:5 ratio by volume and phase separation was performed. The RNA-containing layer was mixed with an equal volume of 100% isopropanol and incubated overnight at −20 °C for precipitation. RNA was pelleted and washed with 80% ethanol and resuspended in nuclease-free water. RNA concentration and purity were determined using a NanoDrop 2000 (Thermo Scientific). RNA for sequencing was sent to University of Massachusetts Medical School Molecular Biology Core Lab for fragment analysis.

The frozen RNA pellets were shipped on dry ice to Genewiz (Germany) for RNA library preparation and sequencing. The RNA sequencing workflow after RNA isolation included initial PolyA selection–based mRNA enrichment, mRNA fragmentation, and random priming with subsequent first- and second-strand complementary DNA (cDNA) synthesis. Afterward, end-repair 5′ phosphorylation and adenine nucleotide (dA)–tailing was performed. Last, adaptor ligation, polymerase chain reaction (PCR) enrichment, and Illumina NovaSeq technology–based sequencing with 2× 150–base pair (bp) read length were carried out. Fastq files were loaded into the DolphinNext platform (https://dolphinnext.umassmed.edu/) and the Bulk RNA sequencing pipeline was used. Fastq files were aligned to both the human (hg19) genome. Once aligned, the files were run through RSEM for normalization. Differential expression analysis was performed using the DEBrowser platform (https://debrowser.umassmed.edu/).

Pathway analysis was performed by combining the Gene ontology analysis (https://geneontology.org/) and TopFunn (https://toppgene.cchmc.org/) results.

### Single Cell RNA-seq

Single-cell library preparation was performed using Chromium™ Single Cell 3’ GEM Library & Gel Bead Kit v3 (Cat# 1000092, 10X Genomics) according to manufacturer’s instruction. The libraries were sequenced on the NovaSeq system (Illumina) using the NovaSeq® 6000 S4 Kits v1.5 (35 cycles; Cat# 20044417, Illumina) performed by Azenta Life Science. The sequencing outputs were processed using the CellRanger software v3.1.0 on the Massachusetts Green High Performance Computer Cluster (GHPCC), Reads were mapped to human reference genome GRCh19 (Ensembl 19). Data analysis was performed using Seurat v4.1.0 within R version 4.0.2 environment(31).

### CD34 Positive Cell Enrichment

CD34^+^ cells were enriched from *de novo* progenitor cells collected from each respective 3D hydrogel tissue explant culture using the EasySep Human CD34 Selection (StemCell Technologies, #17856). HUCB CD34^+^ cells were isolated from whole blood using Lymphoprep (StemCell Technologies, #07801) to collect mononuclear cells then purified using EasySep Human Cord Blood CD34 Selection (StemCell Technologies, #17896). HUCB CD34^+^ cells were then seeded in Matrigel Matrix (Cat# 356231, Corning) for 14-17 days for HSC expansion. After HSC culture expansion, HUCB CD34^+^ cells in Matrigel Matrix were then harvested using Dispase (Cat# 354235, Corning) for 40 minutes followed by additional 14 minutes of Trypsin-EDTA (Cat# 15400-054, Gibco) and Collagenase I (Cat# LS004197, Worthington). HUCB CD34^+^ cells harvested from Matrigel Matrix were subsequently enriched using EasySep Human Cord Blood CD34 Selection (StemCell Technologies, #17896). Enriched CD34^+^ were then either used for FACS, colony forming unit assay, or *in vitro* differentiation.

### Colony Forming Unit Assay and Monocyte/Macrophage In Vitro Differentiation

Enriched CD34^+^ cells were resuspended to a final volume of 1.5 mL with semisolid Methocult (StemCell Technologies, #04404) and then plated in 6-cm dishes and incubated at 37°C in 5% CO_2_. Between 6 and 14 days, the number of colony-forming unit-erythrocyte (CFU-E) and CFU-granulocyte-erythrocyte-monocytes-macrophage (CFU-GEMM), were scored under a microscope. HSC expansion using isolated CD34^+^ cells were cultured in StemSpan (StemCell Technologies, #09645), 1% Pen/Strep, 50ng/ml SCF (PreproTech, # 300-07-10UG), 50ng/ml Flt3 (PreproTech, # 300-19-10UG), 20ng/ml TPO (PreproTech, # 300-18-10UG), 20ng/ml IL-6 (PreproTech, # 200-06-20UG), 0.75 uM StemRegenin 1 (SR1, StemCell Techologies, # 72342), and 35nM UM729 (StemCell Technologies, # 72332). Monocyte differentiation was induced from HSC for 5-7 days in RPMI (Gibco, #11875093), 10% FBS, 1% Pen/Strep, 50ng/mL SCF (PreproTech, # 300-07-10UG), 50ng/ml Flt3 (PrepreoTech, # 300-19-10UG), 20ng/ml TPO (PreproTech, # 300-18-10UG), 25ng/ml M-CSF (PreproTech, # 300-25-10UG). Inflammatory M1 Macrophage differentiation was induced from monocytes for 24-36 hours in in RPMI (Gibco, #11875093), 5% FBS, 1% Pen/Strep, and 100ng/mL IFNγ (PreproTech, # 300-02-20UG). Anti-inflammatory M2 Macrophage differentiation was induced from monocytes for 24-36 hours in in RPMI (Gibco, #11875093), 5% FBS, 1% Pen/Strep, and 10ng/mL IL-4 (PreproTech, # 200-04-5UG) and 10ng/mL IL-13 (PreproTech, # 200-13-2UG).

### Flow Cytometry

Flow cytometry assays were performed on BD LSR. Viable cells were determined with live/dead cell marker (Invitrogen™ eBioscience™ Fixable Viability Dye eFluor™ 455UV, ThermoFisher Scientific #65-0868-14). The following fluorescent conjugated antibodies at a 1:50 dilution were used:

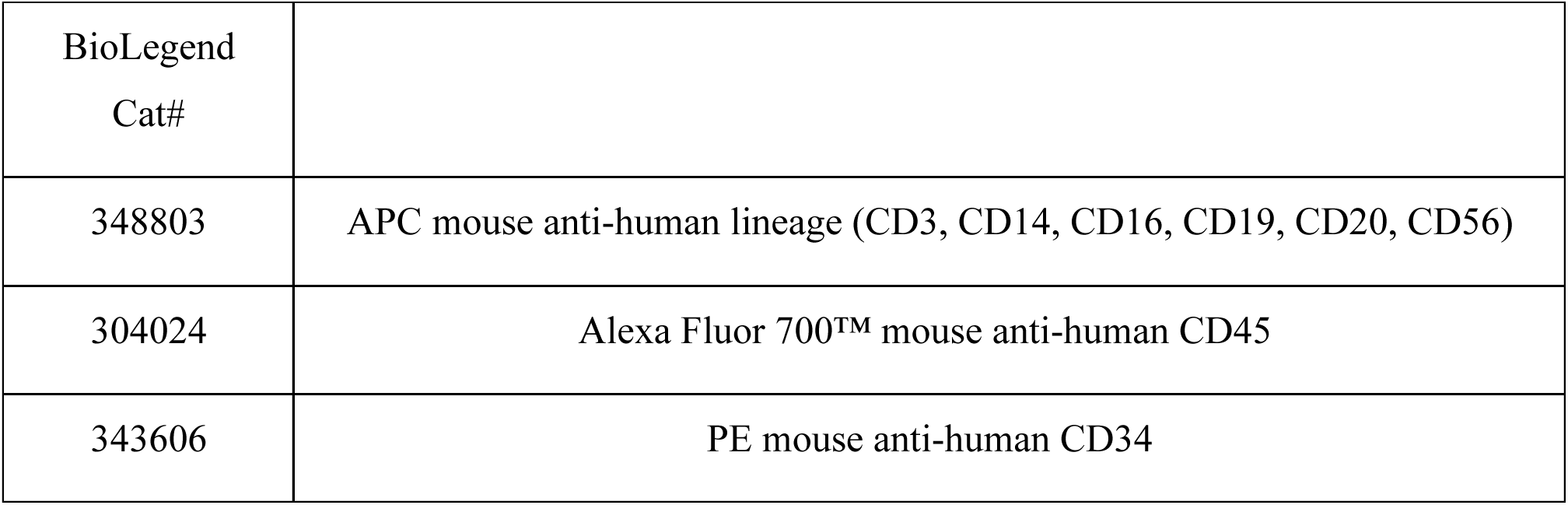
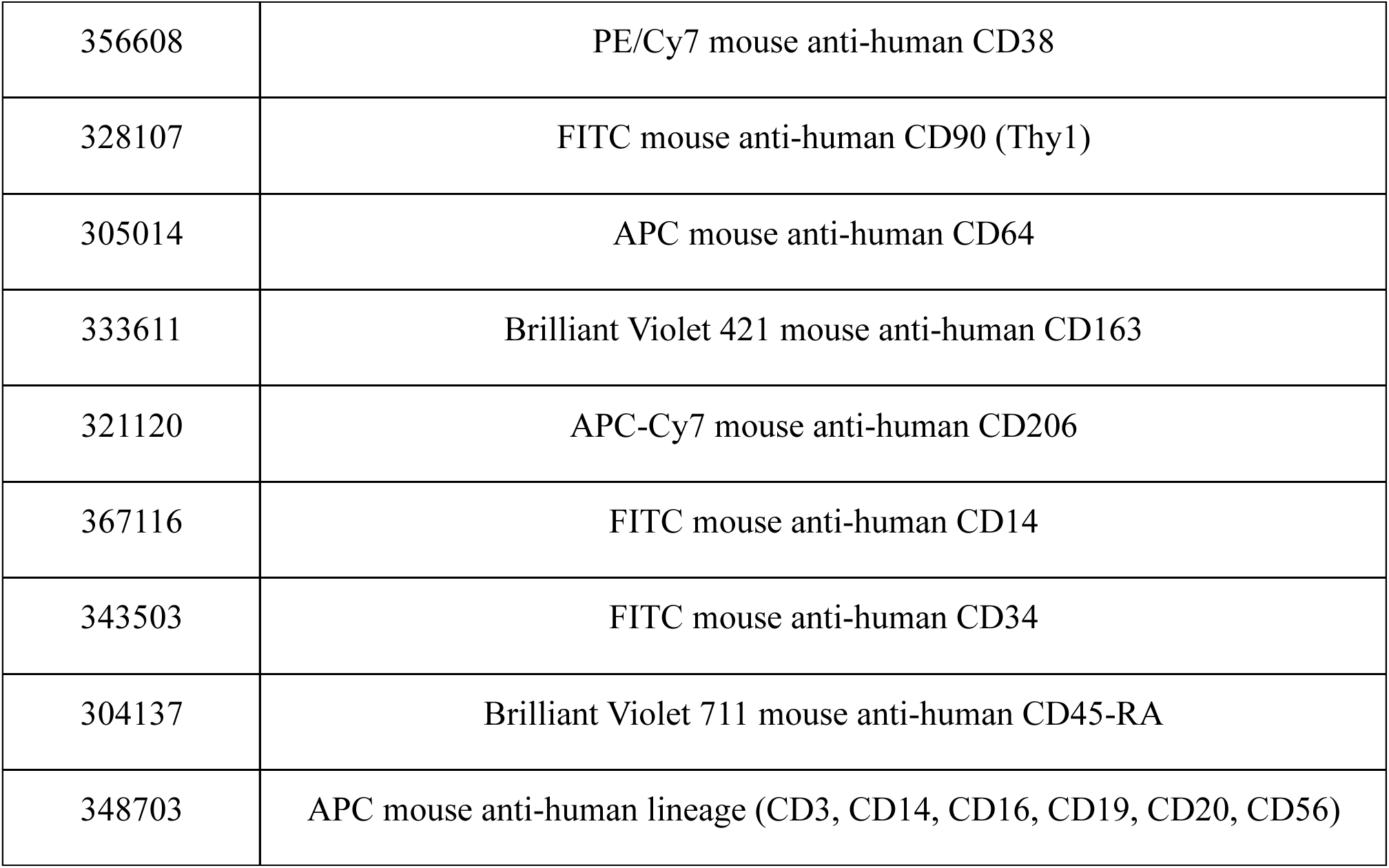

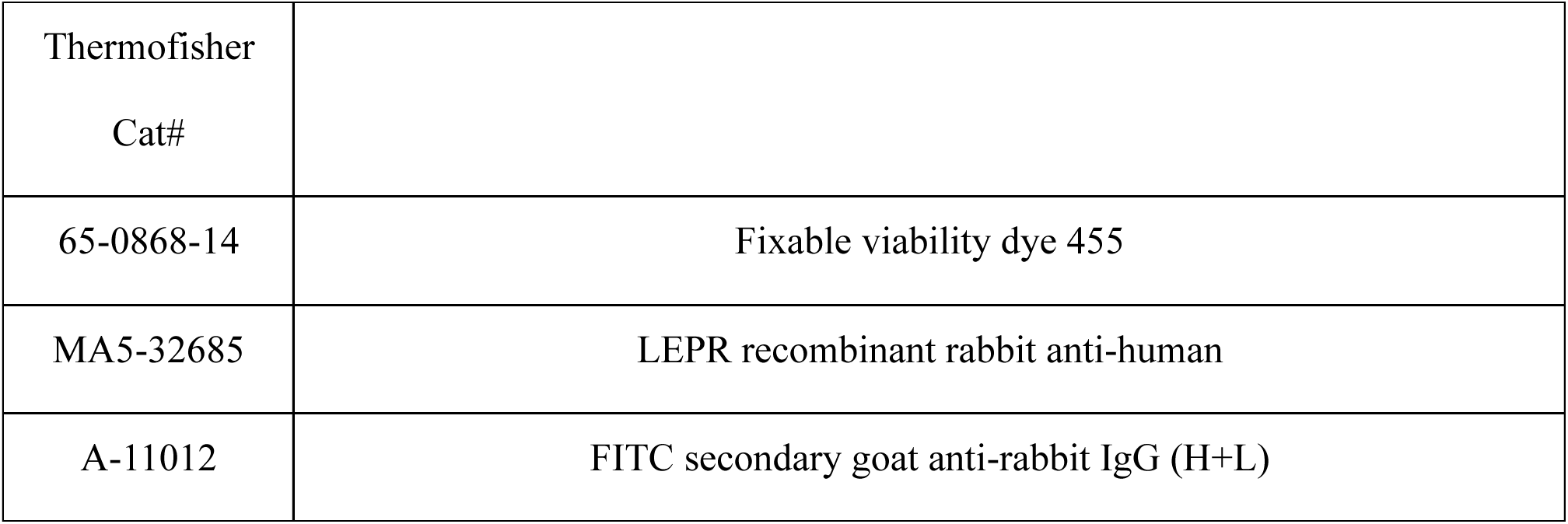

Flow cytometry data analysis was performed with FloJo software (10.10.0).

### HSC-induced Differentiation Towards Monocytes and Macrophages

CD34 positive cells enriched using magnet beads (StemCell) were cultured in complete RPMI 1640 medium (10% FBS, 100 U/ml penicillin, 100 U/ml streptomycin, with 50 ng/ml SCF, 20 ng/ml TPO, and 50 ng/ml Flt3 with 25 ng/ml M-CSF (Peprotech). After 5-7 days, cells were induced to differentiate towards M1 macrophage by changing the medium to the M1 induction medium (complete RPMI 1640 medium, 5% FBS, 100 U/ml penicillin, 100 U/ml streptomycin, and 5 ng/mL IFN-gamma for 24-48 hours. Cells were then collected and stained FACS analysis. Stained cells were run through a BD LSRII or BD FACSymphony A5 flow cytometer and data were analyzed using the FlowJo software (10.10.0).

### Multiplex enzyme-linked immunosorbent assay (ELISA)

Monocytes derived from BMY and BMR organoids (50,000 cells/well in 96-well plate) were stimulated with 100ng/ml of ultra-pure LPS (InvivoGen Catalog #tlrl-3pelps) for 24 hours. Cytokine/chemokines levels were screened in the supernatants by multiplex ELISA by EVE Technologies using their Human Cytokine/Chemokine Panel A 48-Plex Discovery Assay® Array (HD48A).

### Monocytes leptin stimulation

Monocytes differentiated from bone marrow organoid derived CD34 positive cells enriched using the methods described above were cultured in complete RPMI 1640 medium (10% FBS, 100 U/ml penicillin, 100 U/ml streptomycin, with 50 ng/ml SCF, 20 ng/ml TPO, and 50 ng/ml Flt3 with 25 ng/ml M-CSF (Peprotech) with or without 100ng/ml of human recombinant leptin (MilliporeSigma). After 4-8 days, cells were then collected and stained FACS analysis. Stained cells were run through a BD LSRII or BD FACSymphony A5 flow cytometer and data were analyzed using the FlowJo software (10.10.0).

## Author Contributions

TTN: hypothesis generation, conceptual design, experiment design and performance, data analysis, manuscript preparation. ZYL: conceptual design, experiment design and performance, data analysis, manuscript preparation. AS: experiment design and performance. AD: experiment design and performance, data analysis. TD: experiment design and performance, data analysis. HC: experiment design and performance, manuscript preparation. MC: experiment design and performance, manuscript preparation. DK: experiment design and performance, manuscript preparation. SJ: experiment design and performance, data analysis. LK: experiment design and performance, manuscript preparation. JSR: manuscript preparation. RZ: experiment performance, NKC: experiment design and performance. LTT: conceptual design, manuscript preparation. MB: conceptual design, manuscript preparation. LMM: conceptual design, manuscript preparation. KAF: conceptual design, manuscript preparation. EDR: supervision of work, hypothesis generation, conceptual design, manuscript preparation, obtaining funding. SC: supervision of work, hypothesis generation, conceptual design, manuscript preparation.

## Acknowledgements

This study was supported by NIH grants DK089101 and DK123028 to SC, RC2 DK116691 to EDR, Vascular and Endovascular Society Early Career Award, Wylie Scholar Award, and 1K08DK134k52 to TTN. We acknowledge the UMass Chan IT department for computing infrastructure. We acknowledge the use of services from the following UMASS Cores: Flow Cytometry Core, Molecular Biology Core, and Morphology Core. The graphical abstract and flow diagrams were created with BioRender.com.

## Figure Legends

**Supplemental Figure 1.**
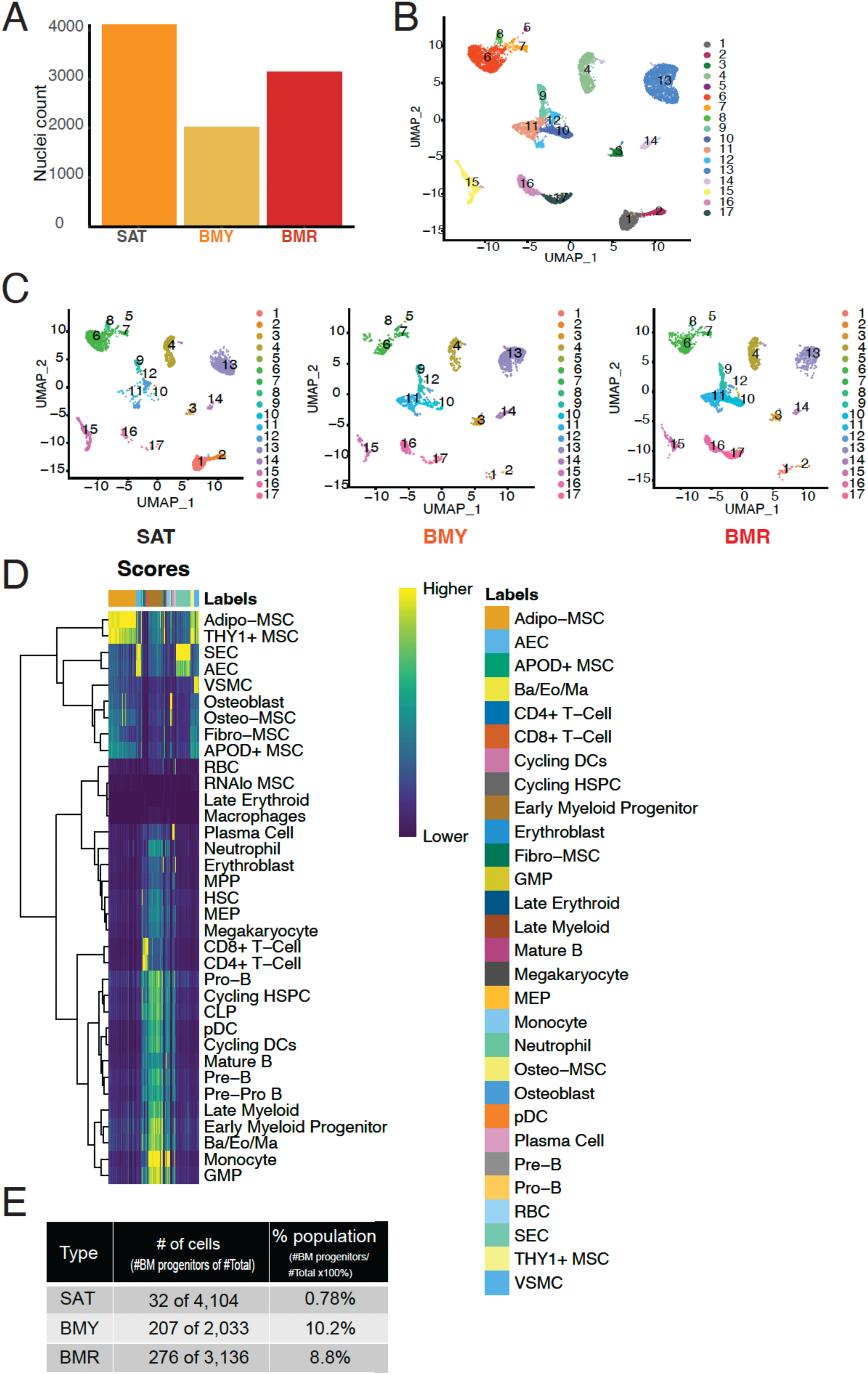
Features of bone marrow and subcutaneous adipose tissue harvested from human donors. (A) Number of nuclei recovered for snRNA Seq. (B) Dimensional reduction plot of snRNA Seq data using UMAP of all cells recovered from SAT, BMY, and BMR tissue depots (n = 1 human donor). (C) Dimensional reduction plot of snRNA Seq data using UMAP for SAT, BMY, and BMR tissue depot (n = 1 human donor). (D) Heatmap of the assignment score for each cell (column) and cell type label (row) from snRNA Seq data of all cells recovered from SAT, BMY, and BMR tissue depots based on Bandyopadhyay *et al.* (E) Number of bone marrow (BM) progenitor cells retrieved from snRNA Seq.

**Supplemental Figure 2.**
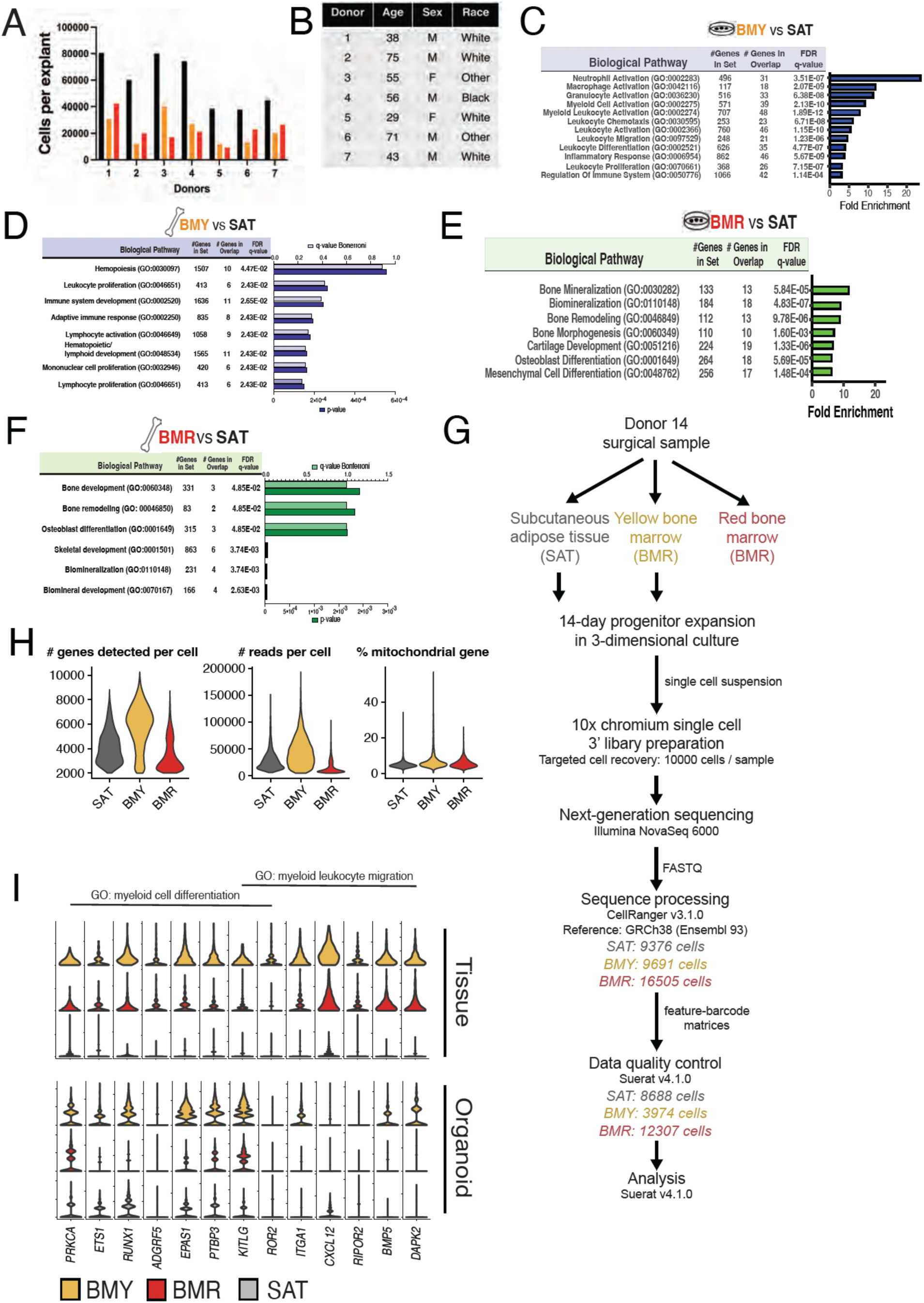
Progenitor cells expanded from bone marrow organoids exhibit a biological pathway enrichment pattern like that of native tissue. (A) Distribution of the total number of sprouting cells per tissue 1mm^3^ tissue explant according to adipose depot type cultured in pro-angiogenic media. (B) Donor demographics for 3D organoid tissue explant culture cultured in mesenchymal stimulating media. (C) Pathway enrichment analysis of differentially expressed genes in organoid cells derived from BMY in comparison to SAT cultured in mesenchymal stimulating media, q-Bonferroni value cut off <0.05 and fold change >4. (D) Pathway enrichment analysis of differentially expressed genes in BMY tissue in comparison to SAT, q-Bonferroni value cut off <0.05 and fold change >4. (E) Pathway enrichment analysis of differentially expressed genes in organoid cells derived from BMR in comparison to SAT cultured in mesenchymal stimulating media, q-Bonferroni value cut off <0.05 and fold change >4. (F) Pathway enrichment analysis of differentially expressed genes in BMR tissue in comparison to SAT, q-Bonferroni value cut off <0.05 and fold change >4. (G) Schematic overview of sample collection for single cell RNA sequencing processing and analysis. (H) Violin plots representing the distribution of number of genes sequenced per cell, reads per cell, and percentage of mitochondrial genes per cell. (I) Violin plots of myeloid pathway gene expression from SAT, BMY, and BMR tissue based on snRNA-seq compared to organoid based on scRNA-seq (n=1 donor per sample type).

**Supplemental Figure 3.**
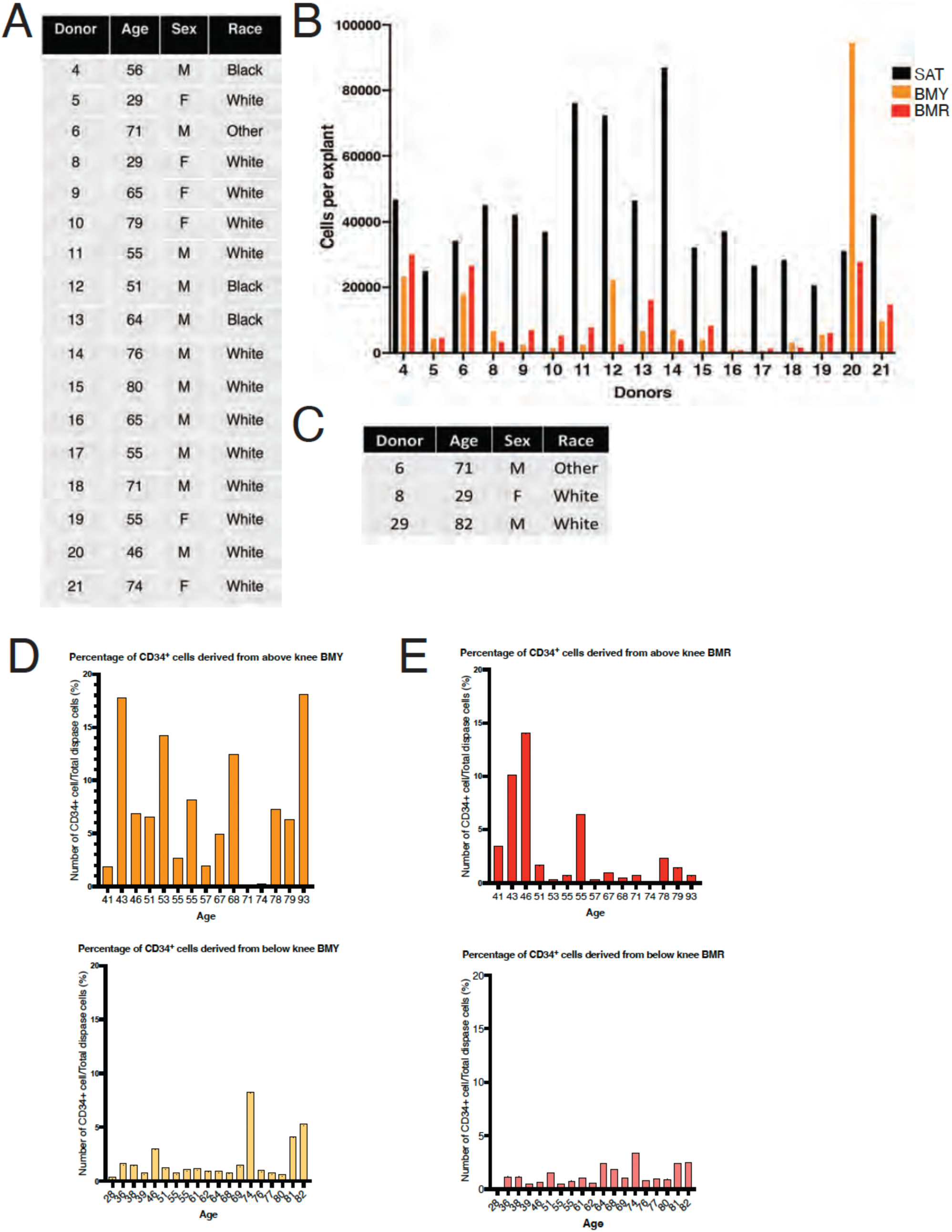
Expansion of progenitor cells from bone marrow tissue and subcutaneous adipose 3D organoid cultured under HSC proliferation conditions. (A) Donor demographics for CD34^+^ cell purification from 3D organoid tissue explant culture cultured in HSC cytokine stimulating media. (B) Distribution of the percentage of CD34^+^ cells purified from total number of sprouting cells per tissue depot type cultured in HSC cytokine stimulating media. (C) Donor demographics for flow cytometry analysis probing for HSPC markers Lin^-^, CD34^+^, CD38^-^, CD90^+^, and CD45^low^. (D) Comparison of the percentage of CD34^+^ cells expanded from total number of sprouting cells per tissue depot harvested from femur above knee BMY and from tibia below knee BMY organoids. The distribution of donor age indicated n = 15 donors. (E) Comparison of the percentage of CD34^+^ cells expanded from total number of sprouting cells per tissue depot harvested from femur above knee BMR and from tibia below knee BMR organoids. The distribution of donor age indicated n = 19 donors.

**Supplemental Figure 4.**
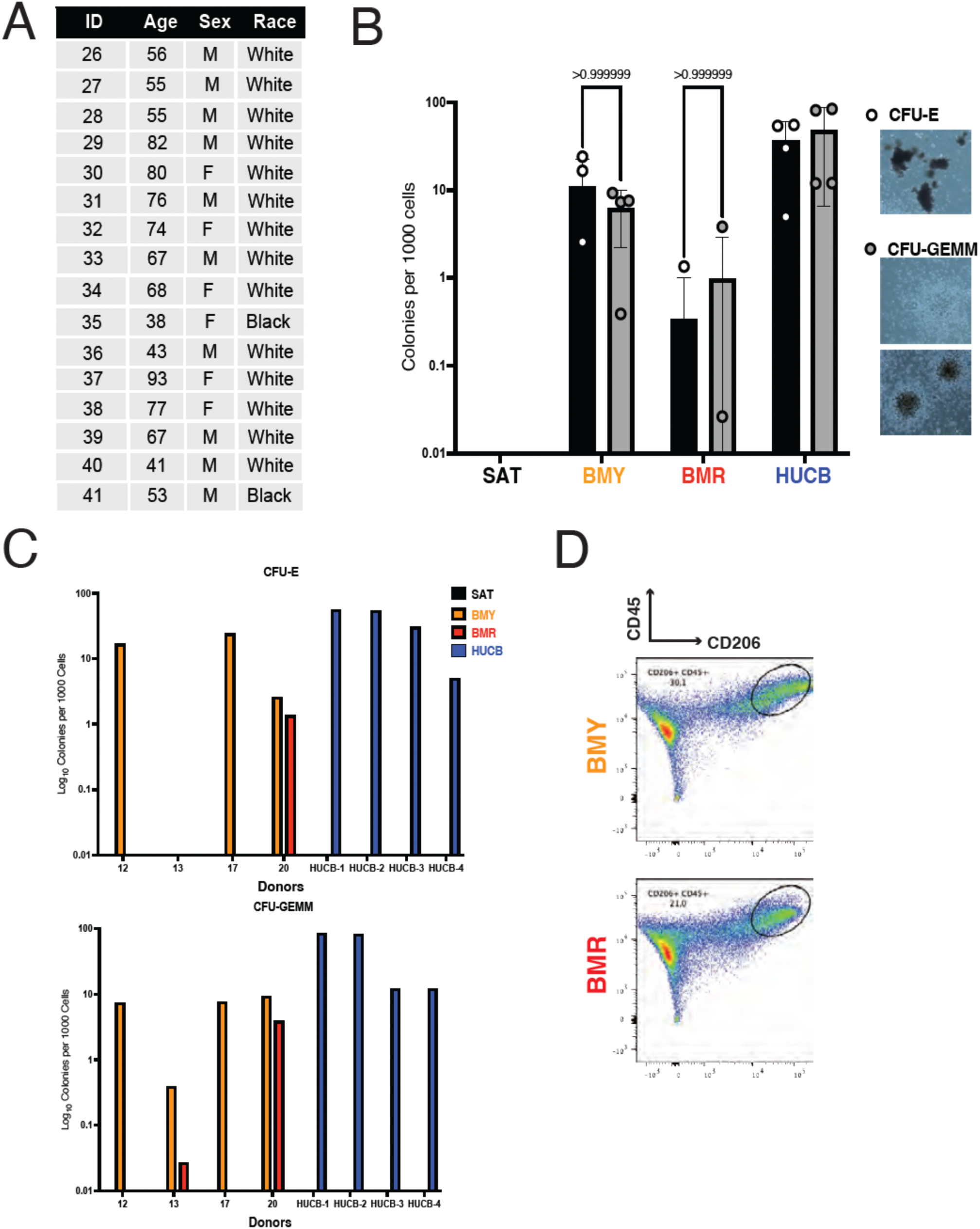
CD34^+^ progenitor cells derived from bone marrow 3D organoid tissue culture are multipotent. (A) Donor demographics for donor samples. (B) Quantification of the mean percentage of colony forming units per 1000 CD34^+^ cells seeded in MethoCult medium. CFU-E and CFU-GEMM colonies were quantified from SAT, BMY, BMR, and HUCB derived CD34^+^ progenitors (n=4 donors). (C) Distribution of the percentage of colony forming units per 1000 CD34^+^ cells seeded in MethoCult medium. Erythroid colonies (CFU-E) and granulocyte, erythroid, monocyte, and macrophage (CFU-GEMM) colonies were quantified from SAT, BMY, BMR, and human umbilical cord blood (HUCB) derived CD34^+^ progenitors (n=4 donors). (D) Representative image of flow cytometry analysis of anti-inflammatory M2 differentiation using CD45 and CD206 cell surface markers.

**Supplemental Figure 5.**
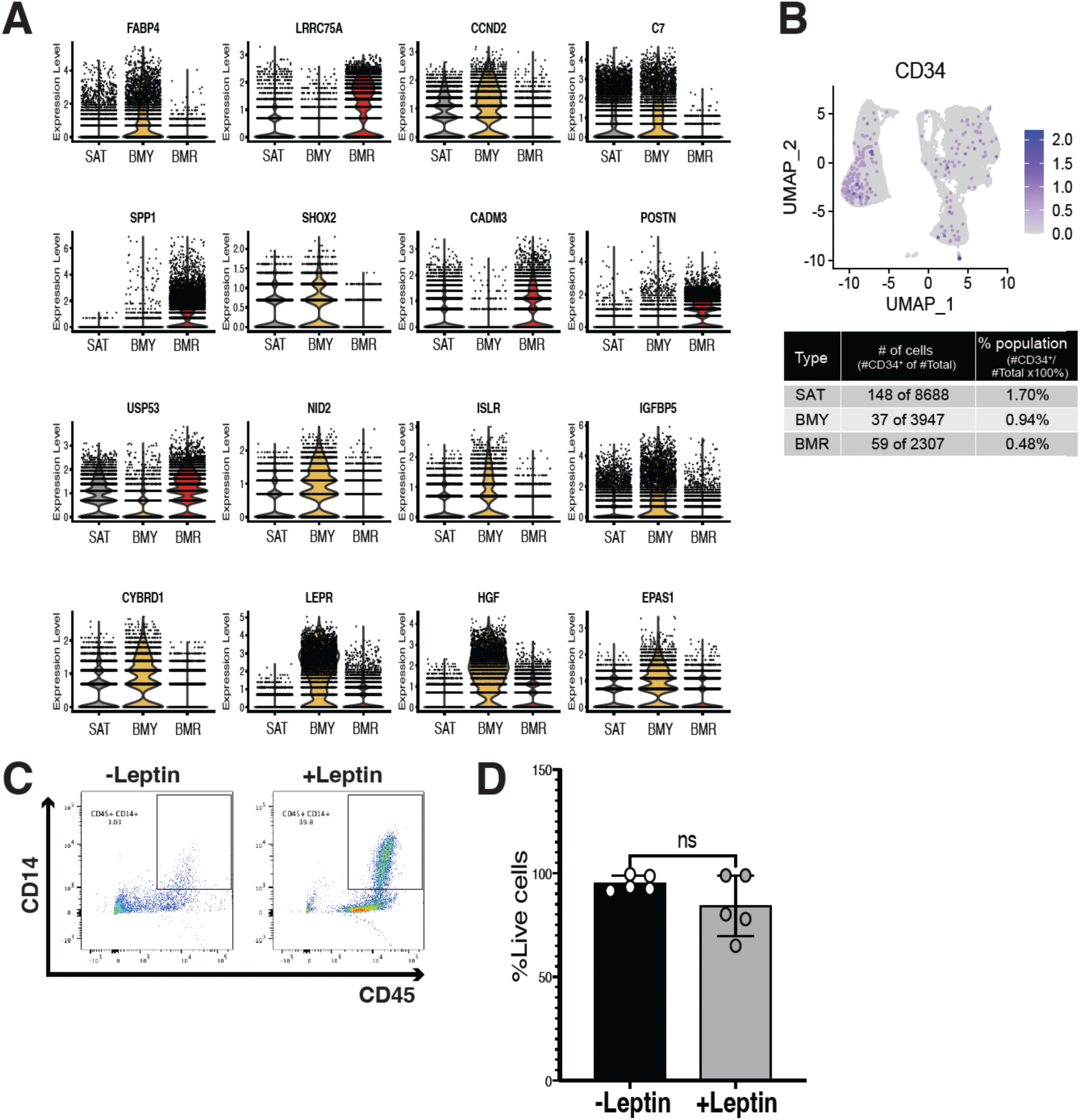
Leptin receptor is differentially expressed in BMY, and leptin stimulation enhances monocyte production. (A) Violin plots of differential expressed genes between SAT, BMY, and BMY organoids (n = 1 donor). (B) Cells expressing CD34 mapped onto the UMAP representation shown in (Figure 2G), with cells expressing the indicated genes highlighted in purple. Number of CD34^+^ cells in SAT, BMY, and BMR tissue depot with corresponding cell population percentage. (C) Representative flow cytometry analysis of CD14^+^ LEPR^+^ cell population cultured with or without exogenous leptin. (D) Quantification of the percentage of live cultured monocytes cultured with or without exogenous leptin based on FACS (n = 5 donors, bars = mean, error bars = SD, ns = no significance).

